# Carboxypeptidase E is efficiently secreted and internalized via lysosomes

**DOI:** 10.1101/2021.06.06.447232

**Authors:** Shir Armoza-Eilat, Michal Caspi, Olga Shomron, Koret Hirschberg, Rina Rosin-Arbesfeld

**Affiliations:** Department of Clinical Microbiology and Immunology, Sackler Faculty of Medicine, Tel Aviv University, Tel Aviv, Israel; Department of pathology, Sackler Faculty of Medicine, Tel Aviv University, Tel Aviv, Israel

**Author notes:** Equal contribution (Koret Hirschberg, Rina Rosin-Arbesfeld).

## Abstract

Carboxypeptidase E (CPE) a key factor in the biosynthesis of most peptide hormones and neuropeptides, is predominantly expressed in endocrine tissues and the nervous system. This highly conserved enzyme cleaves the C-terminal basic residues of the peptide precursors to generate their bioactive form. CPE is a secreted protein; however, the Intracellular pathways leading to its secretion are still obscure. We combined live-cell microscopy and molecular analysis to examine the intracellular distribution and secretion dynamics of fluorescently tagged CPE. CPE was found to be a soluble luminal protein as it traffics from the ER via the Golgi apparatus to lysosomes. Moreover, CPE is efficiently secreted and reinternalized to lysosomes of neighboring cells. The C-terminal amphipathic helix of CPE is essential for its efficient targeting to, and secretion from lysosomes. Fluorescence resonance energy transfer demonstrated that CPE and its substrate neuropeptide Y (NPY) interact in the Golgi apparatus and Immunoprecipitation analysis demonstrated that both CPE and NPY are co-secreted. The implications of the well-defined CPE intra and extracellular routes are discussed.

## INTRODUCTION

CPE is a 35kDa metal-binding exopeptidase that removes one or more basic amino acids (Lys or Arg) from the C-terminal of its substrates (Fricker & Snyder, 1983). CPE is a soluble enzyme that is co-translationally inserted into the ER lumen while its conserved signal sequence is cleaved. The last 25 amino acids at its C-terminal, considered to act as a membrane binding amphiphilic helix (AH) were associated with sorting of substrates into secretory granules via its membrane binding properties (Manser *et al*, 1991). Besides being a biosynthetic enzyme, CPE was shown to have several other functions such as a sorting protein to the regulated secretory pathway, facilitating endocytic and secretory transport processes and regulating signaling pathways (Ji *et al*, 2017; Skalka *et al*, 2016). CPE was shown to expedite the sorting of several of its substrates from the Golgi complex/TGN to the regulated secretory granules. The enzymatic activity sites of CPE are in organelles with pH=5.0 suggesting that it may be active in the Golgi complex and in lysosomes. CPE interacts shortly with its substrates as a protease but may also bind additional proteins for longer time periods to facilitate sorting from the TGN to lysosomes or secretory granules.

Here, we combined biochemical analysis with live-cell microscopy to analyze the intra- and extracellular transport pathways of CPE. Tagged at the C-terminal with a fluorescent protein (FP), CPE was expressed and analyzed in intact living cells, demonstrating that the latter is a soluble luminal protein which is efficiently secreted via lysosomes. The secretion of several truncated CPE-constructs demonstrated that the C-terminal AH is essential for its efficient targeting to and secretion from lysosomes. Fluorescence recovery after photobleaching (FRAP) revealed that CPE is rapidly exported from the ER. Interaction of CPE with its substrate NPY in the Golgi complex was shown using fluorescence resonance energy transfer (FRET) and immunoprecipitation assays. Finally, we found that secreted CPE is reinternalized and apparently concentrated in lysosomes.

## RESULTS

### CPE localizes to the lumen of secretory organelles

CPE is found in secretory granules (Fricker & Snyder, 1983) of tissues that secrete peptide hormones or neurotransmitters (Fricker, 1988). To characterize the subcellular localization of CPE, we examined its distribution in the CPE-expressing PC12 neuroendocrine cell line (Das *et al*, 1992). CPE is detected by immunofluorescence in numerous peripheral and perinuclear punctate structures. The later, colocalize with the trans-Golgi (TGN) marker TGNP-N-10-mCherry (Fig. 1A). Next, we examined the distribution of transfected CPE-GFP in different cell lines (HeLa, HCT116, SW480, HEK293T, and COS7). Figure 1B demonstrates that in all cell lines CPE-GFP is localized to the perinuclear area resembling the Golgi apparatus as well as to the membranes of the endoplasmic reticulum (ER). The identity of the CPE-GFP-positive organelles was verified by live-cell imaging analysis of COS7 cells co-expressing CPE-GFP and the Golgi marker GalT-CFP or the ER marker KDEL-mCherry (Fig. 1C). Co-expression with the cytosolic tubulin marker EB3-mCherry, revealed that the intracellular distribution of CPE-GFP is restricted to the lumen of the cellular organelles. Moreover, co-expression of CPE-GFP and the plasma membrane (PM) marker glycosyl-phosphatidyl-inositol-anchored-mCherry (GPI-mCherry) demonstrated that CPE is absent from the PM (Fig. 1C and supplementary movie S1). Together, the data place CPE in the lumen of secretory organelles without any apparent PM labeling, suggesting that CPE is efficiently secreted.

**Figure 1.**
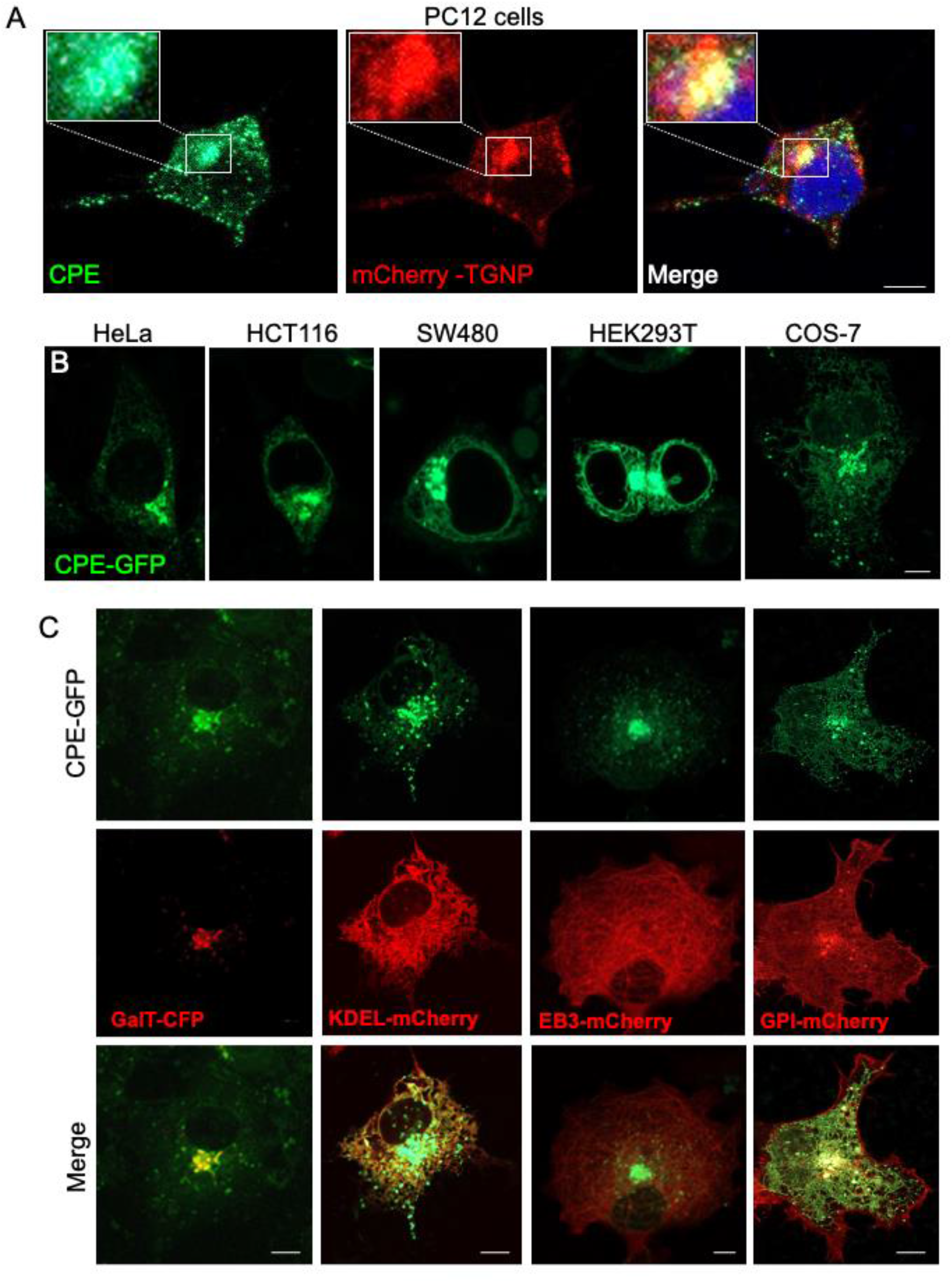
CPE localizes to the lumen of secretory organelles. **A**. Intracellular distribution of endogenous CPE. A image of PC12 cells fixed, permeabilized and stained with DAPI (Blue) and anti-CPE antibody (green), 24 hours after transfection with mCherry-TGNP (red). Scale bar = 5 µm. **B**. Intracellular distribution of CPE-GFP in different cell lines. HeLa, HCT116, SW480, HEK293T, and COS-7 cells were transfected with CPE-GFP. Images of living cells were captured 24 hours later. Scale bar = 10 µm. **C**. Colocalization of CPE with markers of secretory organelles. Confocal images of living COS-7 cells were captured 24 hours after co-transfection of CPE-GFP (green) with GalT-CFP, KDEL-mCherry, EB3-mCherry or GPI-mCherry (red). Scale bar = 5 µm.

### CPE localizes to lysosomes

To further analyze its intracellular distribution, CPE tagged with the three spectral variants mTagBFP, GFP, and mCherry was co-expressed in COS7 cells. Of these three proteins GFP fluorescence is known to be sensitive to endosomal acidic environment. Figure 2A demonstrates that the three CPE-tagged proteins are co-expressed in the Golgi area (Fig. 2A, upper insert). However, the GFP-tagged CPE was excluded from the peripheral punctate structures. This observation suggests that CPE may be expressed in structures such as lysosomes, where the low luminal pH selectively eliminates GFP fluorescence (Cranfill *et al*, 2016; Kneen *et al*, 1998; Shinoda *et al*, 2018). The lysosome marker, LGP120-GFP is an integral membrane protein where the GFP tag faces the cytosol (Bright *et al*, 1997; Fukuda, 1991; Patterson & Lippincott-Schwartz, 2002). Live-cell microscopy demonstrates that CPE and LGP-120 exhibit identical subcellular distribution confirming lysosomal localization of CPE (Fig. 2B). The enlarged panel depicts that CPE occupies the lumen while LGP-120 labels the lysosomal membrane. As a control, we co-expressed LGP-120-GFP with ss-mCherry, a secreted form of mCherry with a cleavable signal sequence (ss) derived from hen egg lysozyme (Patterson et al., 2008). The colocalization between CPE-mCherry and lysoTracker-Blue (which labels the lysosomes) was significantly higher than that of ss-mCherry (Fig. 2C). The association of CPE with lysosomes is also supported by colocalization of the protein with the late-endosome-lysosome marker CD63 (Pols & Klumperman, 2009) (Fig. 2D) as well as with the Rab27a (Fig. 2E), shown to be involved in conventional lysosomal trafficking and exocytosis (Fukuda, 2008). In contrast, neither of the GFP tagged A or B isoforms of Rab11, which represent an early and late endosomal marker respectively, colocalized to any extent with CPE-mTagBFP (Fig. 2 F and G).

**Figure 2.**
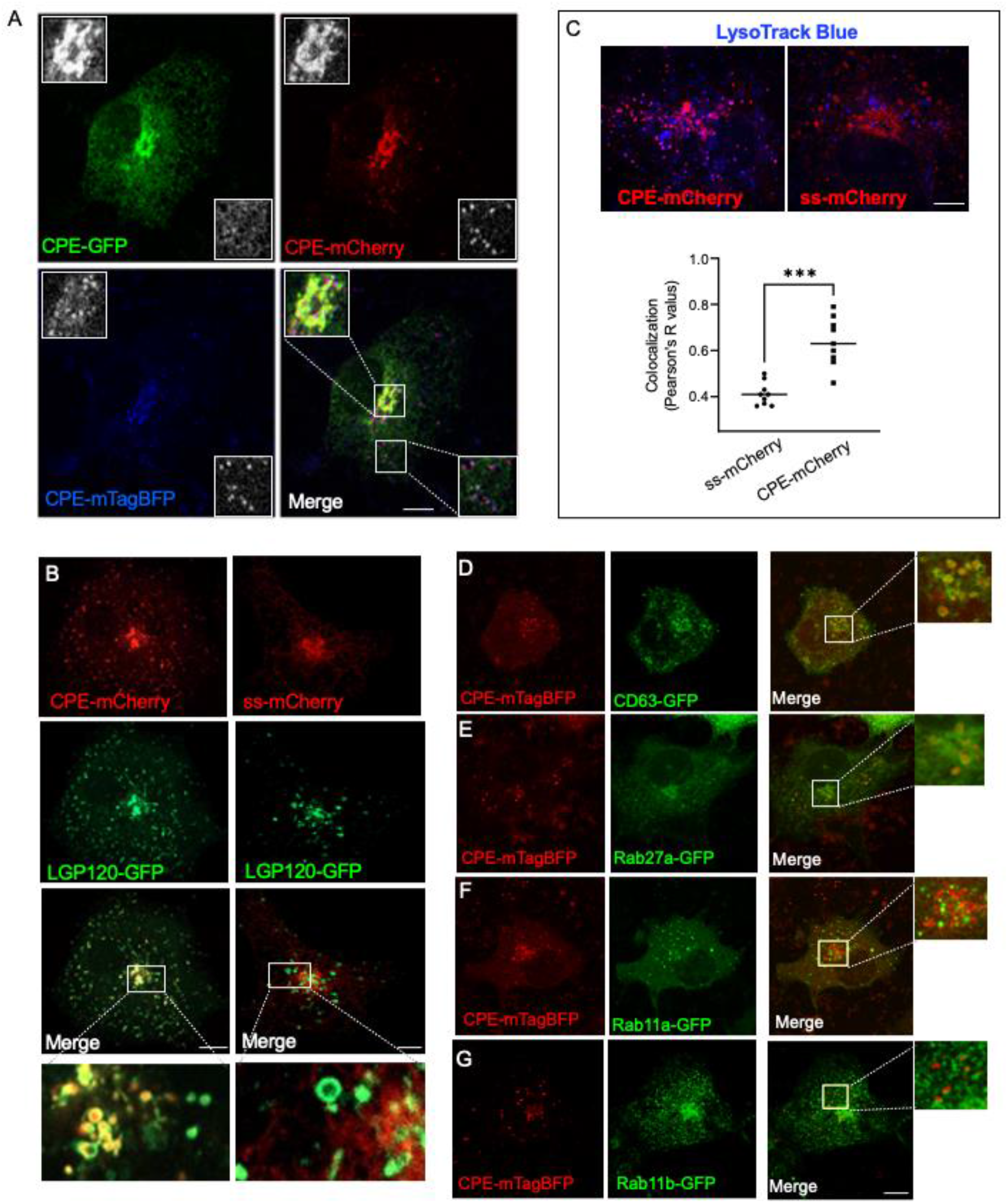
CPE localizes to lysosomes. **A**. CPE localizes to secretory organelles and acidic compartments. COS-7 cells were co-transfected with three spectral variants of GFP: GFP (green), mCherry (red), and mTAG BFP (blue). Image of a living cell was captured 24 hours after transfection. Middle insert showing the Golgi is enlarged x2 and shown at top left side. Bottom insert showing peripheral acidic organelles lacking GFP fluorescence is enlarged X2 and shown at bottom right side. Scale bar = 10 µm. **B**. Comparison of the intracellular distribution of two secreted proteins. Images of living COS-7 cells 24 hours after co-transfection of CPE-mCherry (red, upper panel) or signal sequence (ss)-mCherry (red, lower panel) with the lysosomal marker LGP120-GFP (green) Scale bar = 10 µm. Inserts are enlarged 4x on the right-hand side. Scale bar = 5 µm. **C**. Quantitative analysis of colocalization of CPE-mCherry with LysoTracker. COS-7 cells transfected with CPE-mCherry (red, lower panel) or ss-mCherry (red, upper panel) were labeled with LysoTracker-Blue (Blue) added for 5 min after fixation with formaldehyde. Scale bar = 10 µm. Bottom graph is a quantitative analysis of colocalization using Pearson’s coefficient. The Mann-Whitney t-test was employed; ***P=0.0002. **D**. Colocalization of CPE with endosomal markers. COS-7 cells were co-transfected with CPE-mTAG-BFP (red) and CD63-GFP (green), Rab27a-GFP (green), Rab11a-GFP (green) or Rab11b-GFP (green). Images of living intact cells were captured using a confocal microscope. Scale bars = 10 µm.

### CPE is efficiently secreted

Our results support the hypothesis that CPE-GFP is secreted from COS7 cells but also localizes to the lumen of lysosomes. CPE-GFP is apparently secreted efficiently as it is persistently visualized in the Golgi apparatus and in carriers moving to the cell periphery. When the intracellular distribution of CPE-GFP was compared with that of the secreted ss-mCherry, the two fluorescent proteins did not entirely colocalize, particularly at the cell periphery (Fig. 3A). Cycloheximide treatment of CPE-GFP or ss-mCherry expressing cells, resulted in depletion of both proteins (Fig. 3B, left panels). Additionally, blocking secretory transport with brefeldin A (BFA), an inhibitor of secretory transport (Niu *et al*, 2005) resulted in accumulation of both secreted proteins in the ER (Fig. 3B, right panels). Western blot analysis of CPE-GFP and ss-mCherry detected both proteins in the media of transfected cells (Fig. 3C). Interestingly, the levels of secreted CPE were significantly reduced, but not completely eliminated in the presence of BFA (Fig. 3D), suggesting that CPE may be secreted from a post-Golgi BFA-insensitive route (Laulagnier *et al*, 2011). To further analyze the secretion efficacy of CPE-GFP, we applied fluorescence recovery after photo-bleaching (FRAP) to quantitatively analyze the ER to Golgi transport rates of CPE-GFP and ss-mCherry (Fig. 3E). COS7 cells were co-transfected with both CPE-GFP and ss-mCherry. A region of interest (ROI) over the Golgi was photobleached and images were captured for about 30 min to analyze the ER to Golgi traffic-mediated recovery. Figure 3E demonstrates a typical experiment with two cells in the field of view. The graph shows ROI fluorescence intensity plotted against time and fitted to an exponential equation. The time scale (*1/k*) for Golgi recovery in the two cells examined is 1.35 and 2-fold faster for CPE-GFP compared to ss-mCherry, supporting the notion of a rapid and efficient secretion of the former.

**Figure 3.**
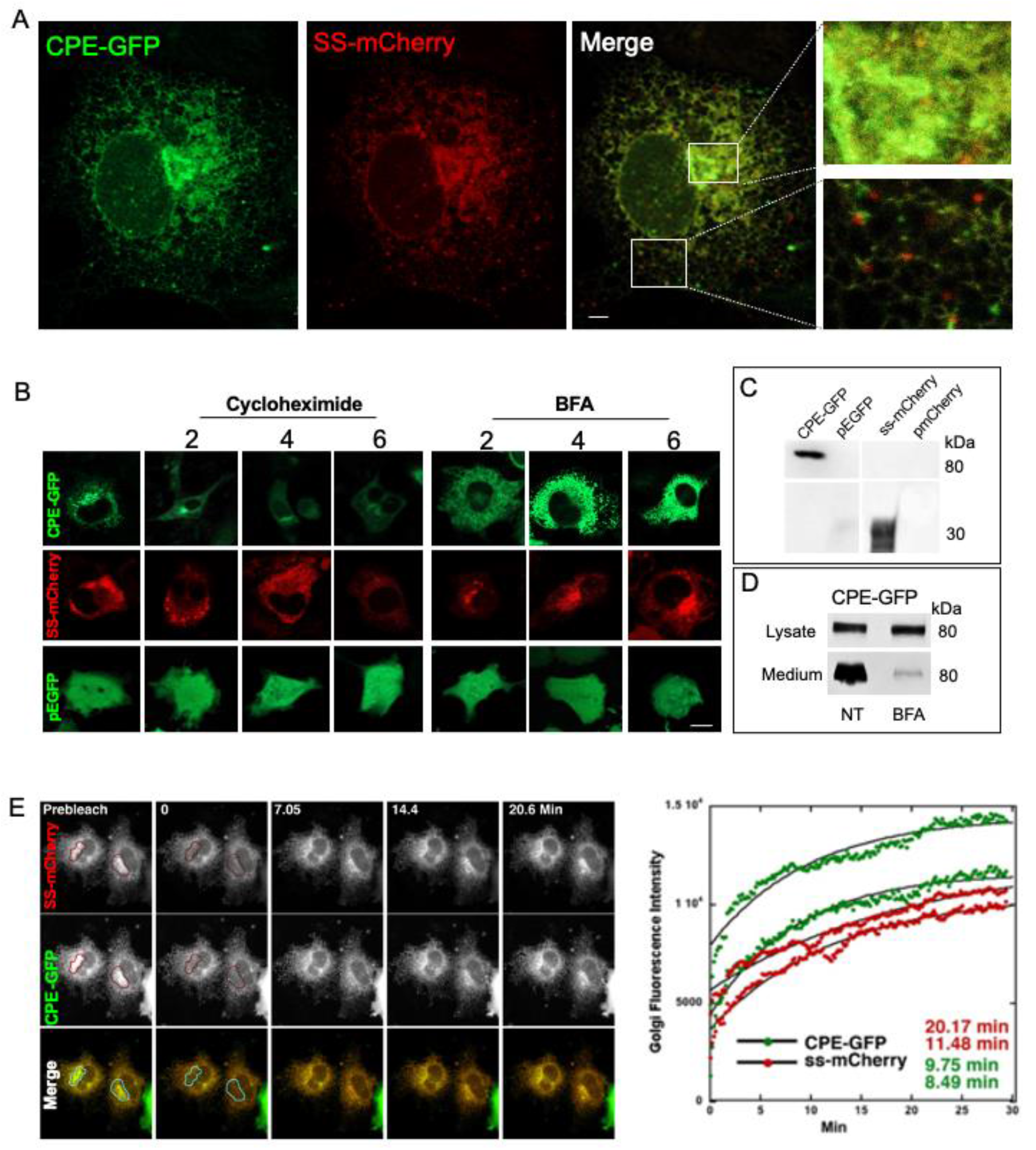
CPE is efficiently secreted. **A**. The secreted proteins CPE-GFP and ss-mCherry do not colocalize at the peripheral punctate structures. Confocal image of a living COS-7 cell co-transfected with CPE-GFP (green) and ss-mCherry (red) 24 hours after transfection. Inserts of the Golgi area and the cell periphery are enlarged 2.5 fold. Scale bars = 10 µm. **B**. Analysis of the effect of Cycloheximide and Brefeldin A on CPE-GFP. COS7 cells transfected with CPE-GFP (top green), ss-mCherry (middle red), or pEGFP (bottom green) were incubated with Cycloheximide (60μg/μl), or Brefeldin A (BFA, 5 μg/μl) for the indicated times. Images of living cells were captured using a confocal microscope. Scale bars = 20 µm. **C**. Western blot analysis of CPE-GFP secretion. The medium of cultured HEK293T cells, transfected with CPE-GFP or SS-mCherry was collected after 24 hours and analyzed on a Western blot using anti-GFP and anti-mCherry antibodies. **D**. Western blot analysis of CPE secretion in the presence of BFA. HEK293T cells transfected with CPE-GFP were incubated with BFA (5 μg/μl) for 4 hours and the cell lysate and medium were analyzed by a Western blot analysis using an anti-GFP antibody. **E**. Quantitative analysis of ER to Golgi trafficking of CPE-GFP and ss-mCherry. COS7 cells were co-transfected with CPE-GFP (middle panel and green) and ss-mCherry (top panel and red). A region of interest over the Golgi (red lines) was photobleached using high power lasers and images were captured for 30 min. Graph at the right-hand side shows the changes in fluorescent intensity of CPE-GFP (green dots) and ss-mCherry (red dots) within the Golgi ROI of the two cells. Black lines were generated by fitting the data to a single exponential equation. The time scales of recovery (values of the inverse of the exponential constant) are indicated. Scale bars = 10 µm.

### CPE is secreted via lysosomes

We and others demonstrated that CPE is found in lysosomes [Fig. 2, supplementary movie 2, and (Saito *et al*, 2011)]. Based on our results with BFA (Fig. 3D) we tested the hypothesis that CPE may be secreted from lysosomes. To this end we analyzed the effects of chloroquine (CQ), an inhibitor of lysosomal acidification and function (Hamano *et al*, 2008) (Fedele & Proud, 2020) on CPE-GFP secretion. CQ was reported to block protein trafficking in and out of lysosomes by raising their intraluminal pH (Andrei *et al*, 1999; Ling *et al*, 1998; Luo *et al*, 2011; Tapper & Sundler, 1990). Moreover, lysosomal secretion of FABP4 was demonstrated using CQ (Villeneuve *et al*, 2018). In the presence of CQ, CPE-GFP accumulated in large peripheral punctate organelles (Fig. 4A). Moreover, WB analysis demonstrated that CQ as well as Bafilomycin A1 (BafA1, a reagent known to disrupt autophagosome-lysosome fusion and lysosome acidification (Mauvezin *et al*, 2015) inhibited CPE-mCherry secretion. In accordance, CPE secretion was enhanced in the presence of ionomycin, a calcium ionophore that triggers calcium release thereby inducing lysosomal secretion (Bennett *et al*, 1979; Carvalho *et al*, 1982). As shown in Figure 4D the secretion of CPE was increased following ionomycin treatment, further supporting the existence of a CPE lysosomal exocytosis pathway. The WB in Fig. S1 demonstrates that the effect of CQ and BafA1 can be reproduced in PC12 cells that secrete endogenous CPE (Fig. S1).

**Figure 4.**
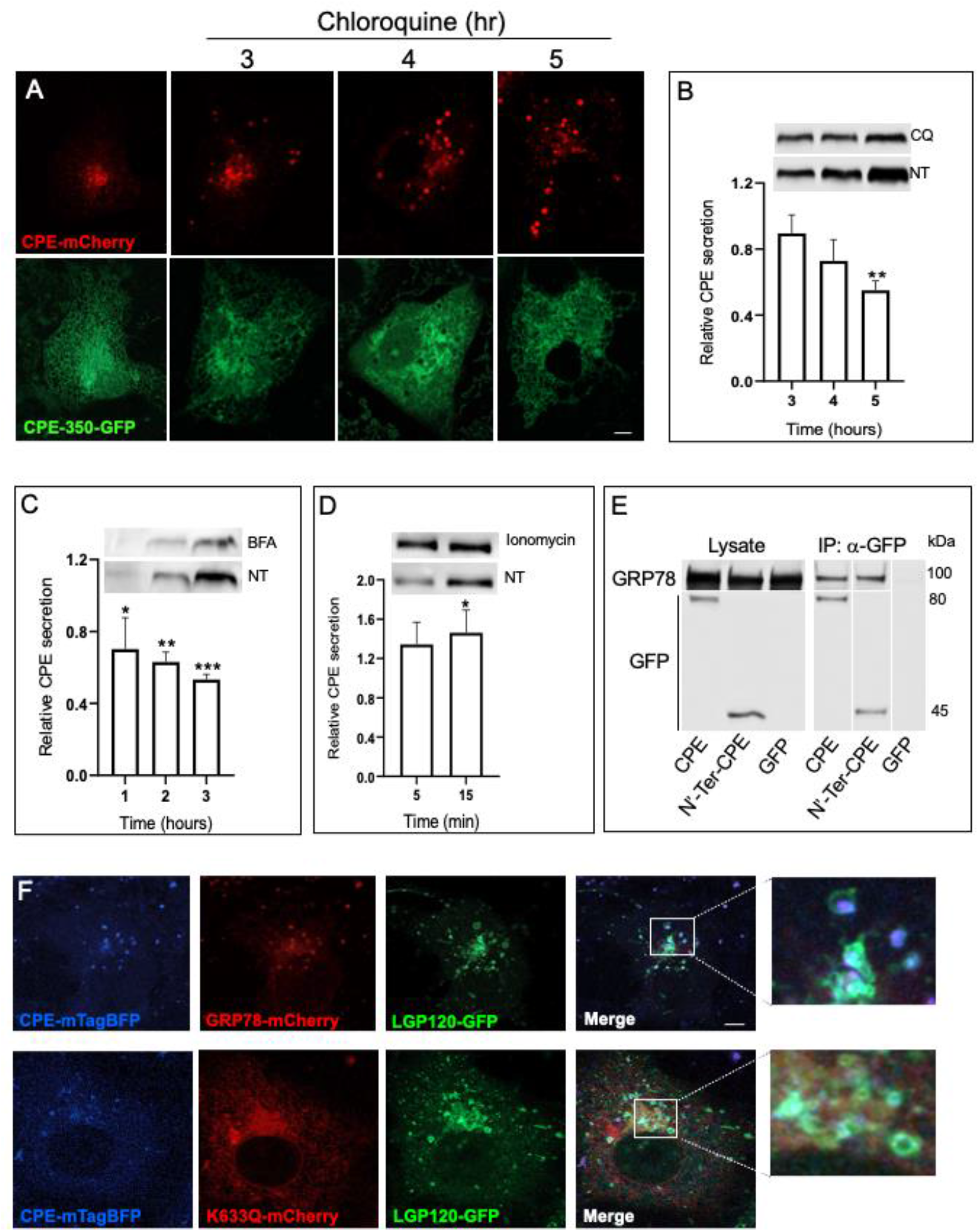
CPE traffics with GRP78 and is secreted from lysosomes. **A**. CPE-mCherry accumulates in lysosomes in the presence of chloroquine. Living intact COS-7 cells were incubated with chloroquine (50 µM) for the indicated times 24 hours after transfection with CPE-mCherry. **B-D**. WB analysis of the effect of chloroquine, Bafilomycin A1, and ionomycin on Lysosomal secretion of CPE-GFP. CPE-GFP secretion was analyzed by WB in HEK293T cells 48 hours post transfection with CPE-GFP in the presence of either 50 µM chloroquine, 100 µM Bafilomycin A1, or 10 µM ionomycin (and 1mM CaCl_2_ in PBS). Equal samples from the cell media were collected at the indicated times, separated by SDS–PAGE and analyzed by WB using an anti-GFP antibody. Quantified band intensities for three separate experiments are shown. Two-way ANOVA test was applied with Sidak’s multiple comparisons test. P=0.006, **P=0.003 (CQ); P < 0.0001, *P = 0.038, **P = 0.0062, ***P = 0.0009 (BafA1); P = 0.027, **P = 0.0167 (ionomycin). **E**. GRP78 coimmunoprecipitates with CPE-GFP. Cell lysates from HEK293T cells expressing either full length CPE-GFP, the N’-terminal #$% of CPE (N’-Ter-CPE-GFP), or EGFP were analyzed 48 hours post transfection. Immunoprecipitation (IP) was performed using an anti-GFP antibody with detection by both anti-GFP and anti-RFP (mCherry) antibodies. **F**. COS7 cells were co-transfected with CPE-mTAG BFP, GRP78-mCherry (upper panel) or GRP78 K633Q-mCherry (lower panel), and LGP120-GFP. Live cell images were recorded 24 h after transfection by confocal microscopy.

Mass spectrophotometric analysis of proteins that co-immunoprecipitated with CPE-GFP, identified GRP78/BIP. GRP78 is a well-characterized secreted ER chaperone that mediates the correct folding and assembly of newly synthesized proteins (Hurtley *et al*, 1989). It was also shown to target aberrant proteins for proteasomal degradation (Corrigall *et al*, 2004; Delpino & Castelli, 2002; Kern *et al*, 2009; Marin-Briggiler *et al*, 2010). Interestingly, a recent study demonstrated trafficking and co-release of GRP78/BIP with β-coronaviruses through lysosomal secretion (Ghosh *et al*, 2020). Thus, we co-expressed CPE-GFP or GFP-tagged CPE N’-terminal domain with GRP78-RFP. Coimmunoprecipitation analysis demonstrated that CPE interacts with GRP78 via its N terminal domain (Fig. 4E). Moreover, the two proteins colocalized in LGP120-GFP positive lysosomes (Fig. 4F, upper panels). Co-expression of CPE-mTag-BFP with an inefficiently secreted GRP78 K633Q mutant (Li *et al*, 2016) significantly reduced targeting of CPE to lysosomes, supporting the premise that GRP78 is involved in the lysosomal secretion of CPE.

### The C’-terminal amphipathic helix is involved in CPE secretion

The 25 C’-terminal amino acids of CPE encode an amphipathic α-helix (AH) domain that has been shown to associate with membranes under acidic pH (5.5–6.5) and plays a role in targeting CPE into the regulated secretory vesicles (Dhanvantari & Loh, 2000; Varlamov & Fricker, 1996). The CPE AH shows high conservation throughout evolution (Fig. S2). It contains of several conserved clusters of alternating acidic and basic amino acids. The secretion of CPE lacking the 33 C’-terminal amino acids was shown to be blocked while CPE that lacks only 23 C’-terminal residues is still secreted (Varlamov & Fricker, 1996). We generated several truncations at the C’-terminal of CPE as shown in Fig. 5A. Contrary to previous reports, we found that CPE lacking the C’-terminal 25 amino acids (CPE-451) is the only construct that gained TGN localization (Fig. 5B). All the other larger truncations were transport incompetent and thus retained in the ER. Interestingly, CPE-451 was detected in lysosomes however to a lesser extent compared to the wildtype CPE (Fig. 5C). WB analysis demonstrated that CPE-451 was also secreted although inefficiently (Fig. 5D). These data strongly support the hypothesis that CPE secretion depends on lysosomal targeting. Moreover, the data render the C’-terminal AH domain as the lysosomal targeting and secretion motif.

**Figure 5.**
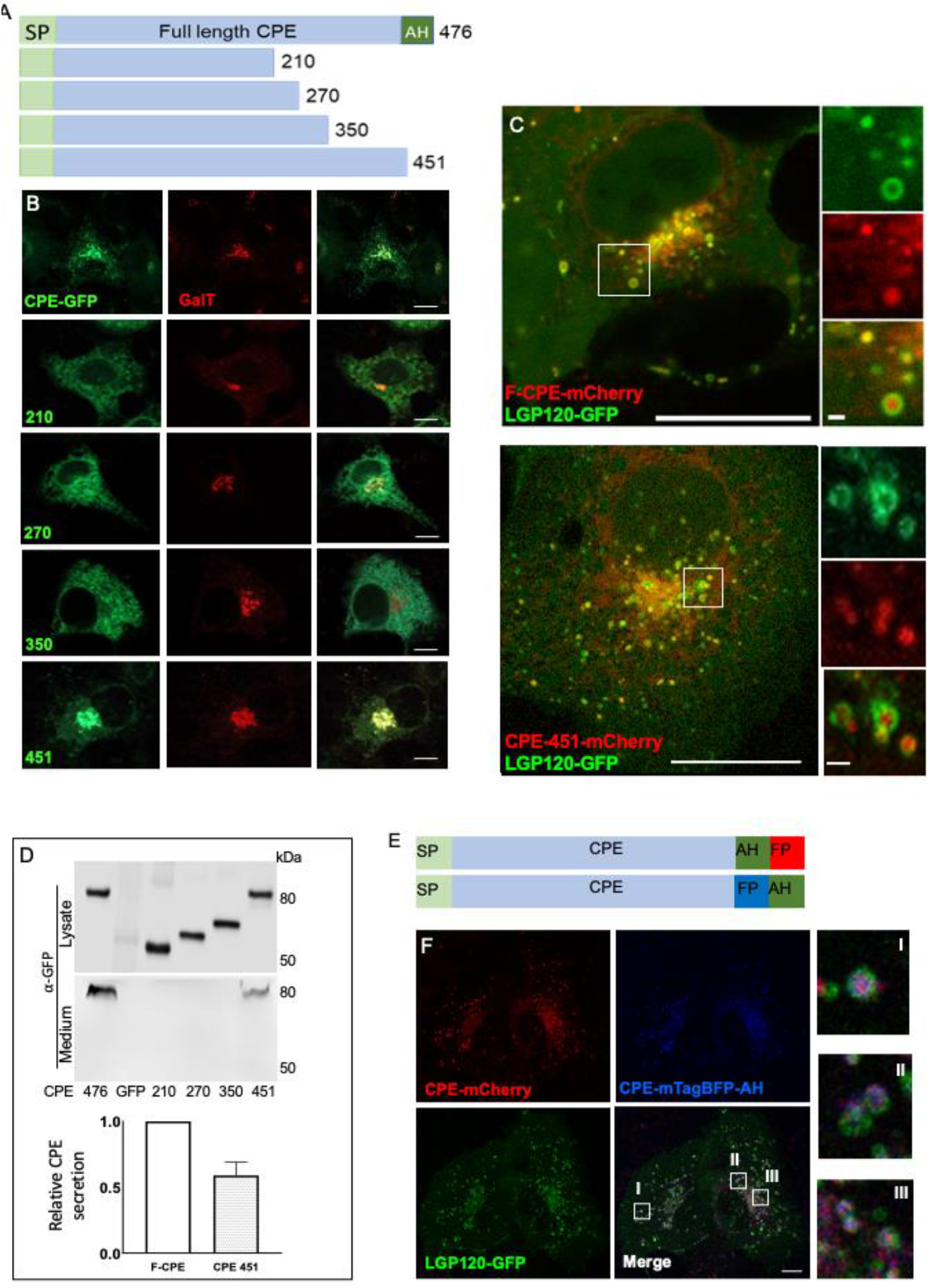
The C’-terminal amphipathic helix of CPE is essential for efficient secretion from lysosomes. **A**. A scheme listing the CPE truncation mutants generated. Light green bars is the cleavable signal peptide. AH - C’-terminal amphipathic helix. **B**. Intracellular localization of the truncation mutants in living cells. COS7 cells were co-transfected with the Golgi marker GalT-CFP (red) and one of the CPE-GFP truncation mutant (green). Confocal Images were obtained 24 hours post transfection. Bar = 10µm. **C**. The CPE_451_-mCherry truncation arrives at lysosomes. COS7 cells were co-transfected with LGP120-GFP (green) and either the full-length CPE-mCherry or CPE_451_-mCherry (red). The images in white square are enlarged 4-fold in images on right hand side. Bars= 10 µm and 1 µm in enlarged inserts. **D**. WB analysis of CPE truncation mutants. HEK293T cells were transfected with one of the CPE truncation mutants. Cells were harvested and culture media were collected 48 hours post transfection. Lysates and media with equal protein concentrations or volumes, respectively, were loaded onto SDS–PAGE gels for WB analysis using an anti-GFP antibody. Graph shows band intensity for secreted wildtype and CPE_451_-GFP. **E**. Scheme of CPE constructs with the FP upstream or downstream to the AH. **F**. The FP at the C’-terminal does not affect CPE localization via steric hindrance on the AH. COS7 cells were co-transfected with LGP120-GFP (green), CPE-mCherry (red), CPE-mTAG-BFP with the AH at the C’-terminus (blue). Images of living cells were captured 24 hours after transfection. Inserts I, II and III are enlarged 5-fold on the right-hand side. Bar = 10 µm.

Our experiments rely on CPE tagged with a fluorescent protein at the C’-terminal. To eliminate the possibility of steric hindrance of the AH motif by the fluorescent protein we re-cloned CPE with the AH positioned downstream to the fluorescent protein (CPE-mTagBFP-AH, Fig. 5E). Live cell fluorescent microscopy of CPE-mTagBFP-AH co-expressed with CPE-mCherry and LGP-120-GFP showed no indication of any effect of the fluorescent protein on CPE intracellular distribution (Fig. 5E).

### Secreted CPE is endocytosed and delivered to the lysosomes but not to the Golgi

Previously it has been reported that a chimeric protein containing CPE C’-terminal 25 amino acids could be recycled back to the TGN from the PM (Arnaoutova *et al*, 2003).The mechanism involved interaction with lipid rafts as well as with ARF6. In our hands, we could not find any evidence for membrane interaction of CPE as it consistently localized to the lumen of organelles. However, we tested whether secreted CPE can be internalized back into the cell. To this end, COS7 cells transfected with CPE-mCherry or ss-mCherry as a control, were re-plated together with cells expressing the lysosomal marker LGP120-GFP (Fig. 6A). Figure 6B shows images captured 24h after re-plating the cells. Secreted CPE-mCherry was internalized by the LGP120-GFP expressing cells and localized to lysosomes (Fig. 6B). A similar experiment, where the acceptor cells expressed the Golgi marker GalT-CFP demonstrated that CPE internalization apparently did not occur via the Golgi apparatus. To verify the results, we repeated the experiment by adding only media from the CPE-mCherry secreting cells (Fig. 6D). Here CPE-mCherry was also detected in lysosomes after 24h as shown in Fig. 6E. These data demonstrate, for the first time to our knowledge that CPE secreted from lysosomes into the media can re-enter and localize to lysosomes of neighboring cells. Thus, the intra and extra cellular routes of CPE remarkably resemble that of corona viruses that are internalized and exit their host cell via lysosomes (Ghosh *et al*., 2020).

**Figure 6.**
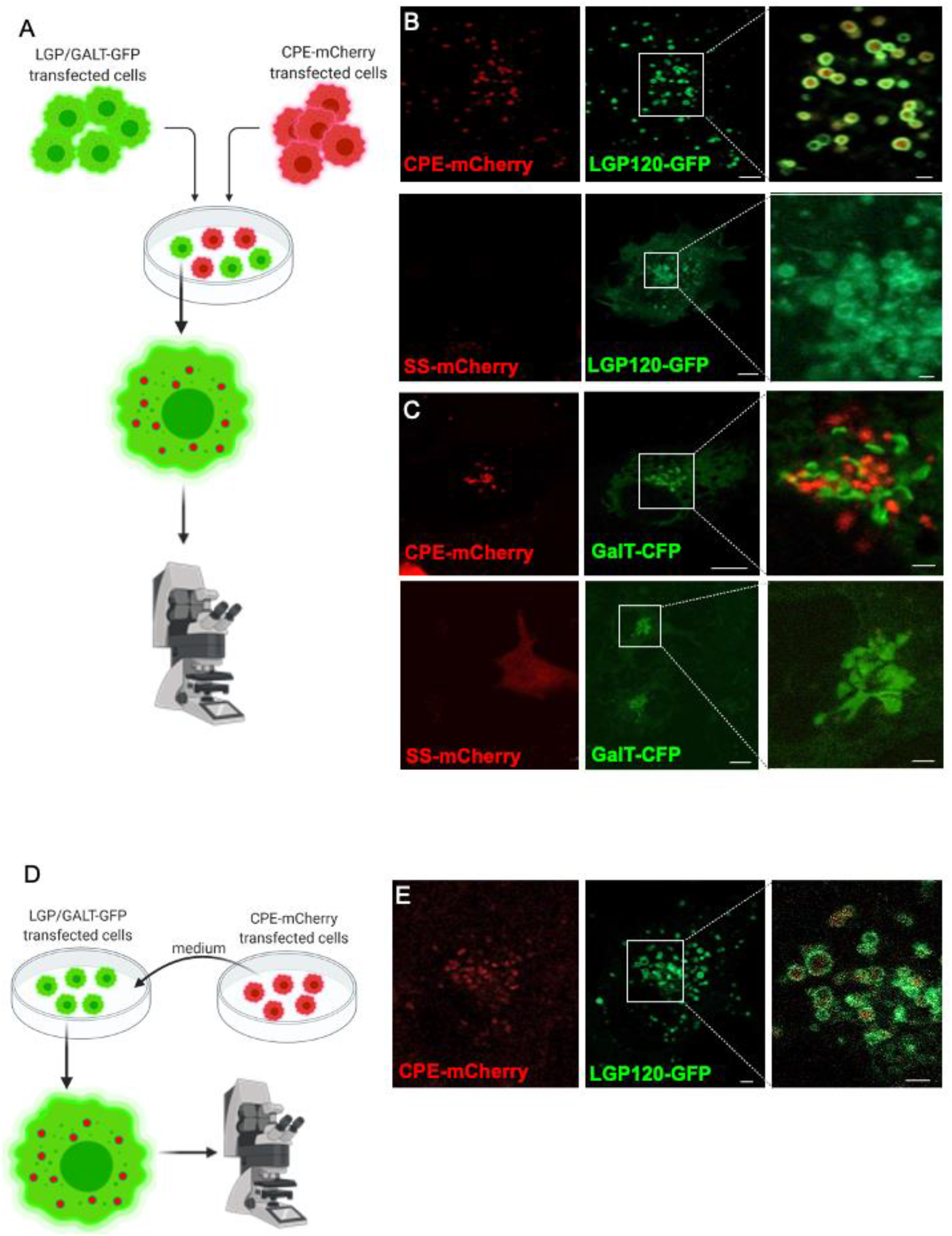
Secreted CPE is endocytosed, delivered, and concentrated in lysosomes but not in the Golgi. **A**. Schematic illustration of the experimental procedure. Cells were cultured together after separately transfecting each with LGP120-GFP or GalT-GFP, CPE-mCherry, or ss-mCherry as a control. Images were captured 24 hours after co-culturing of the cells. **B**. Lysosomal internalization of secreted CPE-mCherry. Images of the cells expressing LGP120-GFP (green) were captured 24 hours after coculturing with the CPE-mCherry or ss-mCherry (red) expressing cells (red). Inserts are regions containing lysosomes magnified 4-fold. Bar = 2 µm **C**. CPE is not internalized to the Golgi apparatus. Same as in B accept that GalT-GFP was used instead of CPE-GFP. Bar = 2 µm. **D**. Schematic illustration of the experimental procedure where only the media from cells transfected with CPE-mCherry was added to cells transfected with LGP-120-GFP. **E**. COS-7 cells transfected with LGP120-GFP (green) were incubated for 30 minutes with media obtained from CPE-mCherry (red) transfected cells. Confocal live cell images are shown. Insert showing lysosomes is magnified 3-fold. Bar = 2 µm.

### CPE and NPY interact and co-localize in lysosomes

Next, we investigated the interaction of CPE with its substrates along its intracellular route. Neuropeptide Y (NPY) is a conserved 36 amino acid peptide, expressed in the peripheral and central nervous systems and is a known substrate of CPE (Brakch *et al*, 1997; Fricker, 1988; Tang *et al*, 2009). Primarily we tested the colocalization of CPE with its NPY substrate. Co-expression of CPE-mTagBFP, NPY-mRFP and LGP-120-GFP demonstrated that both CPE and NPY localized to the lumen of the Golgi apparatus (not shown) and lysosomes (Fig. 7A). Next, we asked to determine if CPE interacts with its NPY substrate. To this end, we used stepwise acceptor photobleaching (Yaffe *et al*, 2015) to analyze FRET interaction between CPE-GFP and NPY-mRFP. The analysis was carried out in the Golgi as the CPE-GFP fluorescence was eliminated by the acidic environment of the lysosomes. Cells expressing both CPE-GFP and NPY-mRFP serving as donor and acceptor, respectively, were fixed as described in the “Methods” section prior to a series of photobleaching of an ROI over the Golgi apparatus. The images in Fig. 7B shows the diminishing fluorescence of the acceptor (upper panel) as well as the resulting increase in donor fluorescence (pseudo-color, lower panel). The FRET efficiency is plotted with the acceptor fluorescence and is compared with a negative control where the donor is LGP-120-GFP. The fluorescent proteins in this control pair were separated by the membrane as the GFP tagged to LGP-120 donor is cytosolic. Thus, the increase in donor fluorescence as a result of photobleaching the acceptor, is indicative of the close proximity of CPE and NPY and their potential interaction. These results were confirmed by immunoprecipitation analysis of CPE-GFP and NPY-mRFP. Moreover, the interaction of the secreted proteins apparently persisted in the medium (Fig. 7C). Figure 7D demonstrates that the transport incompetent CPE-350 truncation did not affect the lysosomal targeting of NYP-mRFP.

**Figure 7.**
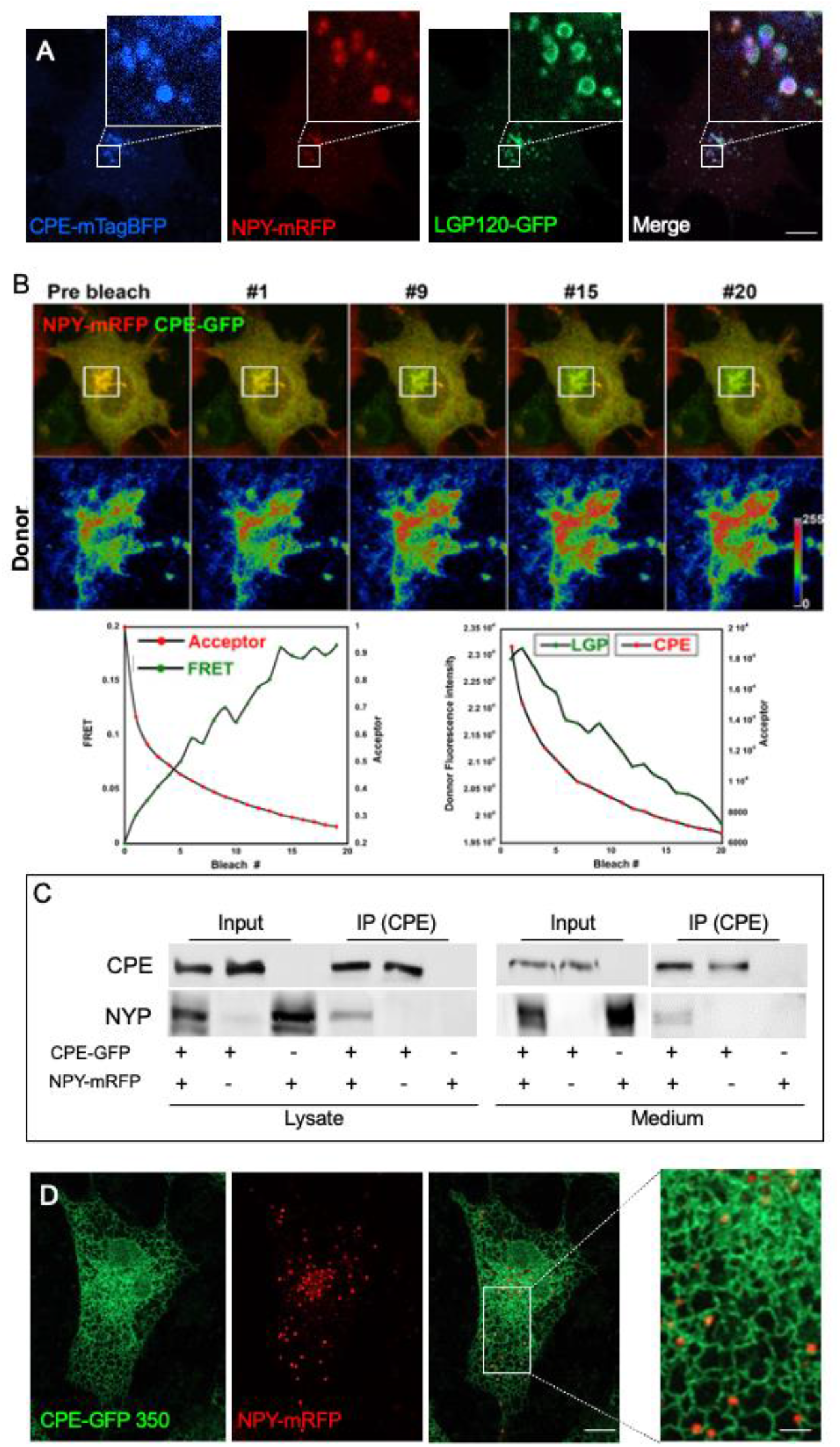
CPE and its substrate NPY interact and co-localize in lysosomes. **A**. CPE and its substrate NPY colocalize to the lumen of lysosomes. COS-7 cells were co-transfected with CPE-mTAG-BFP (blue), NPY-mRFP (red), and LGP120-GFP (green). Images were captured 24 hours post transfection. Inserts are enlarged 5 -fold. Bar = 5µm **B**. FRET analysis demonstrates interaction of CPE and NPY. Cells were co-transfected with CPE-GFP and NPY-mCherry. Stepwise acceptor photobleaching was carried out after fixing the cells with 4% formaldehyde. Top panel shows a typical cell during the experiment. White square is the photobleached region over the Golgi. White square are magnified 4-fold and the green FRET donor channel is shown in pseudo-color with a look up table on the right hand side. Graphs show donor (green) and acceptor (red) fluorescence for the experiment (left) and for a negative control (right)where LGP-120-GFP was used as a donor. **C**. IP analysis confirms interaction of CPE and NPY. HEK293T cells were co-transfected with CPE-GFP and NPY-mRFP. Collected medium and harvested cells were harvested 48 hours post transfection and subjected to IP using anti-CPE antibody. Precipitants were separated on SDS-PAGE and analyzed with anti-GFP and anti-RFP antibodies to recognize the CPE and NYP, respectively. **D**. NPY does not require CPE for its lysosomal targeting. Confocal image of a typical cell co-expressing CPE_350_-GFP and NPY-mRFP captured 24 h after transfection. Insert is enlarged 3-fold. Bar = 10 µm.

Taken together, combining live cell and molecular analysis we delineate for the first time the secretory route of CPE, a key enzyme in the biosynthetic pathway of numerous hormones and neuropeptides. CPE is secreted from- and is internalized directly to lysosomes.

## DISCUSSION

CPE was reported to function in the TGN and dense core secretory granules (Dikeakos & Reudelhuber, 2007; Kim *et al*, 2006; McGirr *et al*, 2013; Ramamoorthy & Whim, 2008; Topalidou *et al*, 2020).The results presented here indicate that CPE may be targeted to, and function in the acidic lumen of lysosomes. The majority of studies describing the secretion of CPE have been based on biochemical analyzes and fractionation assays. Here, for the first time to our knowledge, live cell microscopy combined with biochemical molecular analysis of CPE secretion was carried out. Primarily, we found that counter to previous reports, CPE is a soluble luminal protein that is efficiently targeted to lysosomes from where it is secreted. Direct secretion from the Golgi is demonstrated as well. These findings are in agreement with some reports (Fricker & Snyder, 1983) and contradict others such as those reporting a role for lipid rafts and the small GTPase ARF6 in its targeting (Arnaoutova *et al*., 2003). Moreover, we found that the C’-terminal AH, previously reported to mediate membrane anchoring and retention in the cell, is essential for both efficient lysosomal targeting and secretion. A possible interpretation of the quantitative FRAP analysis is that the AH serves as a sorting signal for CPE which recruits the secretory machinery for efficient transport. The ER to Golgi transport-mediated recovery is significantly faster compared to that of the ss-mCherry. Furthermore, the results of the FRAP experiment are an underestimation when considering a rapid Golgi export of CPE. The deletion of the AH resulted in a significant decrease in CPE secretion. However, its lysosomal targeting was at least in part preserved. The secretion of CPE via lysosomes is compatible with secretion from the regulated secretory pathway found in professional secreting cells. This pathway is regulated by calcium and thus can be activated using reagents such as ionomycin. The role of the co-transported GRP78 in CPE trafficking is not clear. However, GRP78 seems to be obligatory for lysosomal targeting. GRP78 was reported to travel with corona virus proteins that enter and exit cells from lysosomes (Ghosh *et al*., 2020). We hypothesize that the role of GRP78 in CPE trafficking may be to prevent its denaturation or even degradation in the acidic hostile environment of lysosomes. The finding of the re-internalization of secreted CPE to lysosomes is intriguing as CPE was reported to be internalized to the Golgi apparatus. It is yet to be determined whether CPE internalization is specific or rather mediated by bulk flow fluid phase endocytosis. The latter is less likely as the fluorescence of re-internalized CPE was easily visualized in lysosomes whereas in the cell media it was highly diluted to bellow detection levels. The role of CPE in the processing of NPY was described over two decades ago (Brakch *et al*., 1997). The presence of both CPE and NPY in secretory and lysosomal organelles, their interaction in the Golgi apparatus, and their co-secretion, are in agreement with their reported colocalization in dense-core granules of cortical astrocytes (Cheng *et al*, 2013). The interpretation of our FRET data demonstrates a direct interaction between CPE and its NPY substrate. As CPE was also reported to act as a lysosomal targeting chaperon for substrates (Saito *et al*., 2011) it is not clear whether NPY binding results in its processing or simply a persistent interaction aimed to facilitate their targeting to lysosomes. However, based on our results and previous studies, NPY does not require CPE to arrive at lysosomes.

To summarize, we demonstrate that CPE is an efficiently secreted soluble enzyme that is sorted to lysosomes from where it is secreted. The physiological significance of re-internalization of secreted CPE back to lysosomes is yet to be discovered. CPE is a key enzyme that processes numerous hormones and neuropeptides. Thus, it is possible that CPE re-internalization to lysosomes of neighboring cells allows efficient processing of its substrates in acidic organelles by equally spreading the enzyme activity within the entire secreting organ.

## Materials & Methods

### Cell culture, transfection

The PC12 cell line was maintained in RPMI-1640 Medium supplemented with 10% heat-inactivated horse serum, 5% fetal bovine serum, and 100 units/ml penicillin, at 37°C in a humidified incubator with 5% CO_2_. All other cell lines were maintained in Dulbecco’s modified Eagle’s medium (DMEM) supplemented with 10% fetal bovine serum and 100 units/ml penicillin at 37°C in a humidified incubator with 5% CO_2_. Transfection of HEK293T and COS-7 cells was performed using the DNA transfection reagents jetPEI (Polyplus Transfection, Illkirch, France) following the manufacturer’s protocols. The HeLa, HCT116, and SW480 cell lines were transiently transfected using the DNA transfection reagent Lipofectamine 2000 (Invitrogen) following the manufacturer’s protocols.

### Plasmids and reagents

CPE-GFP (pEGFP-N3) was kindly provided by Dr YP Loh (NICHD, NIH, Bethesda, MD, USA). N’-Ter-CPE and the other truncated CPE constructs were constructed by amplifying CPE cDNAs by PCR and subcloning into pEGFP-N3 (Clontech, Oxon, UK) using XhoI and BamHI restriction sites. EB3-mCherry, GPI-mCherry, KDEL-mCherry, GAL T-CFP, Signal Sequence-mCherry (ss-mCherry) was described previously (Dayel *et al*, 1999). NPY-mRFP, CD63-GFP, Rab27a-GFP, Rab11a-GFP, and Rab11b-GFP were kindly provided by Prop. Ronit Sagi-Eisenberg (Department of Cell and Developmental Biology, Sackler Faculty of Medicine, Tel Aviv University). F-CPE-mCherry was constructed using the Gibson Chew Back and Anneal Assembly technique to subclone CPE into pmCherry-N1 (Clontech) vector using XhoI and BamHI restriction sites. LGP120-GFP was purchased from Addgene. CPE-mTAG BFP was constructed using the Gibson Chew Back and Anneal Assembly technique to subclone mTAGBFP into F-CPE GFP vector using BamHI and NotI restriction sites. CPE-mTAGBFP-Amphipathic Helix was constructed by Gibson assembly by sub-cloning CPE without the Amphipathic Helix, mTAGBFP, and the CPE Amphipathic Helix motif into CPE mCherry vector using BamHI and NotI restriction sites. CPE 451 mCherry constructed by Gibson assembly by sub-cloning the 451 amino acids of CPE (without the Amphipathic Helix) into pmCherry-N1 (Clontech) vector using XhoI and BamHI restriction sites.As control vectors we used pEGFP-N3 and pmCherry-N1.All constructs were sequenced to confirm the fidelity of the process. Brefeldin A, Cycloheximide, Chloroquine, BafA1 and Ionomycin were purchased from Sigma-Aldrich (Rehovot, Israel).

### Antibodies

The following antibodies were used: mouse anti-GFP (IB: 1:1,000; IP: 1: 400, Santa Cruz Biotechnology). Rabbit anti-RFP (IB: 1:1000, MBL Life Science, Japan), mouse anti-CPE (IB: 1:1000, IF: 1:200, IP: 1:400, BD Transduction Laboratories, Franklin Lakes, NJ, USA). Anti-mouse and anti-rabbit secondary antibodies were obtained from Jackson Immuno Research (West Grove, PA, USA) (1:10,000). Alexa green (1:500; Molecular Probes, Grand Island, NY, USA) was used for immunofluorescence (IF).

### Confocal laser scanning microscopy, Live cell imaging

COS-7 cells were grown in live-cell imaging chambers (New Biotechnology Ltd, Jerusalem, Israel). Twenty-four hours post transfection, cells were imaged in DMEM by a Zeiss LSM Pascal confocal laser scanning microscope (Carl Zeiss MicroImaging, Jena, Germany). Fluorescence emissions resulting from 405 nm excitation for CFP, 488 nm excitation for GFP, and 543 nm excitation for mCherry were detected using filter sets supplied by the manufacturer. The confocal and time-lapse images were captured using a Plan-Apochromat ×63 1.4-numerical-aperture (NA) objective (Carl Zeiss MicroImaging). The temperature on the microscope stage was held stable (37°C) during time-lapse sessions using an electronic temperature-controlled airstream incubator. Images and movies were generated and analyzed using the Zeiss LSM Zen software and NIH Image and ImageJ software (W. Rasband, National Institutes of Health, Bethesda, MD, USA).

### Fluorescence recovery after photobleaching (FRAP)

For FRAP measurements, a ×63 1.4-NA Plan-Apochromat objective was used on an inverted LSM800 system. Photobleaching of GFP and mCherry was performed using four to six rapid scans with the laser at full power. A region of interest (ROI) over the Golgi was photobleached at both 488 and 565 nm using a high-power laser, and images were captured for about 30 min to record the ER to Golgi traffic-mediated recovery. ROI fluorescence intensity was plotted against time. Data was fitted to a single exponential equation:

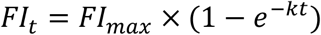

*FI*_*max*_ *=* max fluorescence after recovery, *FI*_*t*_ = fluorescence at time <u>t</u>, and *k* = exponential constant in min^-1^.

Pre- and postbleach images were captured at 0.5 to 3 s intervals, using low laser intensity. Fluorescence recovery in the bleached region during the time series was quantified using LSM Zen software (Carl Zeiss MicroImaging). For presentation purposes, 16-bit confocal images were exported as TIFFs, and their contrast and brightness were optimized in Adobe Photoshop software (San Jose, CA) or in ImageJ (Wayne Rasband, NIH, Bethesda MD).

### FRET analysis

For acceptor photobleaching FRET, cells were grown on glass coverslips. The cells were fixed by addition of paraformaldehyde to the medium to a final concentration of 4% and incubation for 15 min at room temperature. The cells were then washed twice with PBS containing 1% FCS, and once with PBS, and were mounted for visualization on the microscope. The mCherry-tagged acceptor was photobleached in a ROI over the Golgi complex. The 488 nm or 514 nm laser line was used to assess the acceptor photobleaching. FRET efficiency (E) was calculated from the CFP-channel images according to: E_*F*_ = (F_post_-F_re_)/F_post_, where F is the intensity of GFP fluorescence using the 488 nm laser before (pre-bleach) and after (post-bleach) photobleaching of mCherry using a high-power 543 nm laser.

### Immunofluorescence (IF)

PC12 and COS-7 cell lines grown on coverslips, were washed in PBS 24 hours following transfection, and fixed in PBS containing 3.7% paraformaldehyde for 20 min. After two washes with PBS, the fixed cells were permeabilized with PBS containing 0.1% Triton X-100 (PBT) for 10 min and blocked with PBT containing 1% BSA (BBT) for 1 h. Subsequently, cells were incubated in BBT at room temperature with primary and secondary antibodies for 60 and 30 min, respectively. Finally, cells were incubated for 5 min with 10 μg/ml 4’,6-Diamidino-2-phenylindole (DAPI; Sigma-Aldrich) to stain cell nuclei, washed with PBS, and mounted with Fluoro-Gel mounting medium (Electron 31 Microscopy Sciences). GFP, mCherry and mTagBFP were detected without staining. For LysoTracker assay COS-7 cells was grown on imaging chambers (New Biotechnology Ltd, Jerusalem, Israel). Twenty-four hours post transfection LysoTracker was added for 5 min at 37°C and the cells were fixed in PBS containing 3.7% paraformaldehyde for 20 min. Cells were visualized by a confocal microscopy.

### CPE internalization assay

In order to evaluate internalization of CPE into cells, COS-7 cells were transfected with the indicated plasmids in a 6-well plate, and then, 24 h post transfection, the cells were combined and re-plated. Twenty-four hours later, the pooled cells were visualized by a confocal microscope. In some experiments, 24 h post transfection, the medium of LGP120-GFP transfected cells was replaced with medium from COS-7 cells transfected with F-CPE-mCherry or ss-mCherry. The cells were incubated for 30 min at 37°C followed by visualization with a confocal laser microscope.

### SDS-PAGE and Western blot analysis

Forty-eight hours following transfection, medium was collected and centrifuged at 11,000g for 5 min. The cells were then washed with PBS and solubilized in lysis buffer (100 mM NaCl, 50 mM Tris, pH7.5, 1% Triton X-100, 2 mM EDTA) containing protease inhibitor cocktail (Sigma-Aldrich). Cell lysates were incubated on ice for 20 min, then were homogenized and clarified by centrifugation at 14,000 g for 15 min at 4°C. The protein concentration was determined by Bradford reagent. Samples with equal amounts of protein were separated on 10% SDS–polyacrylamide gel electrophoresis (SDS-PAGE), and the proteins were then transferred onto nitrocellulose membranes. The membranes were blocked with 5% low-fat milk and incubated with specific primary antibodies at 4°C (as indicated), washed with PBS containing 0.001% Tween-20 (PBST), and incubated with the appropriate horseradish peroxidase-conjugated secondary antibody for 1 hour at room temperature. After washing in PBST, membranes were subjected to enhanced chemiluminescence (ECL) detection analysis.

### Statistical analysis

Data were analyzed using GraphPad Prism software (version 9.0, GraphPad, La Jolla, CA) and are presented as the mean with standard deviation. Statistical analysis of variance (ANOVA) was performed: P < 0.05 was regarded as statistically significant.

## Supplementary figures

**Figure S1.**
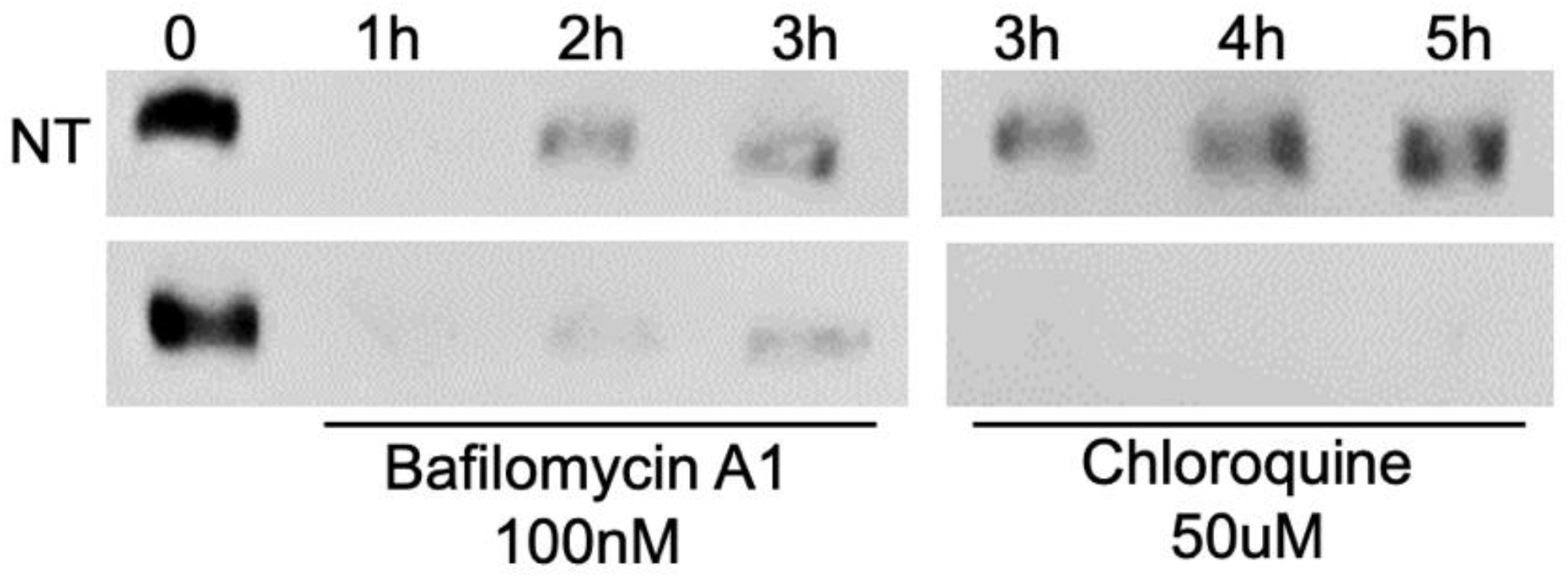
Endogenous CPE is secreted from lysosomes in PC12 cells. WB analysis of CPE secretion by PC12 cells. Cells were grown to confluency and at time 0 were incubated in the absence or presence of BAFA1 or chloroquine for the indicated times. obtaining sample 0. Media samples were analyzed by WB using anti-CPE antibody.

**Figure S2.**
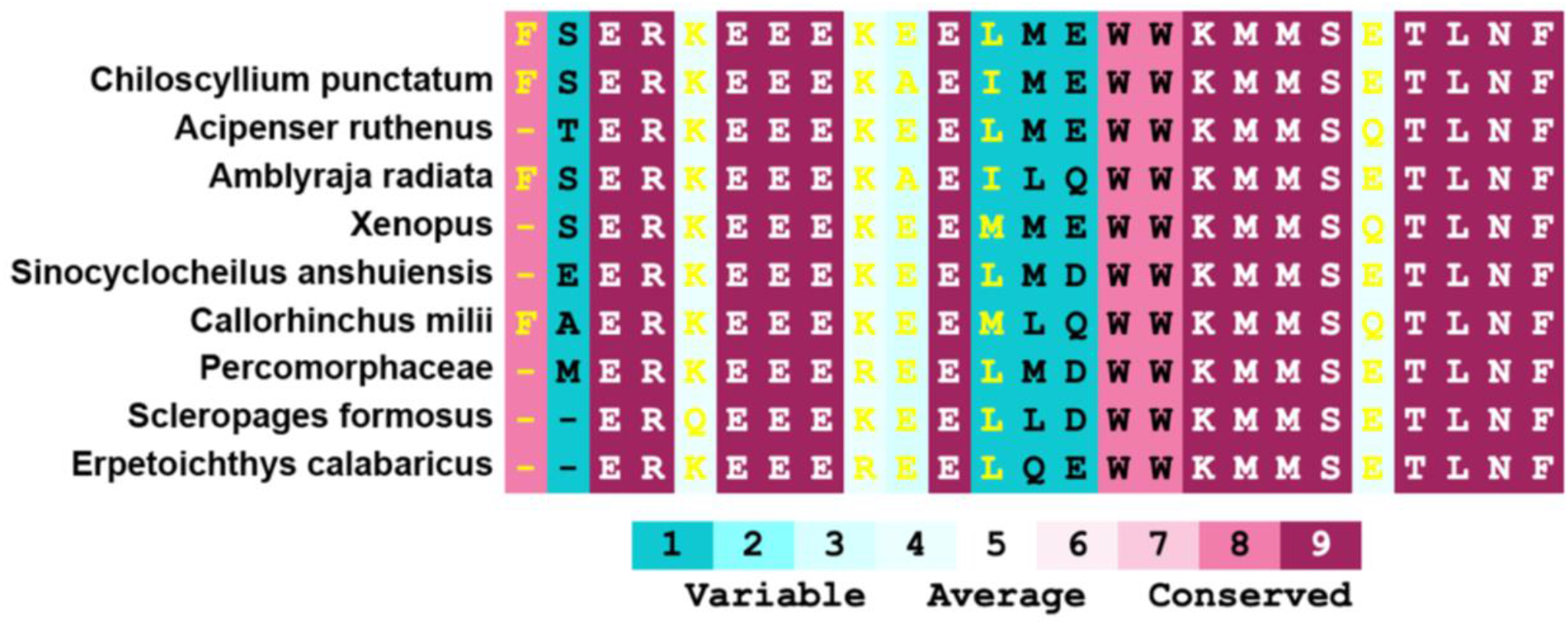
The CPE AH is highly conserved throughout evolution. Human CPE C’-terminal AH was analyzed using ConSurf (Ashkenazy *et al*, 2016) alignment of mouse CPE AH (top) with primitive vertebrates

## Movie caption, Double click image to watch movie

**Movie S1.**
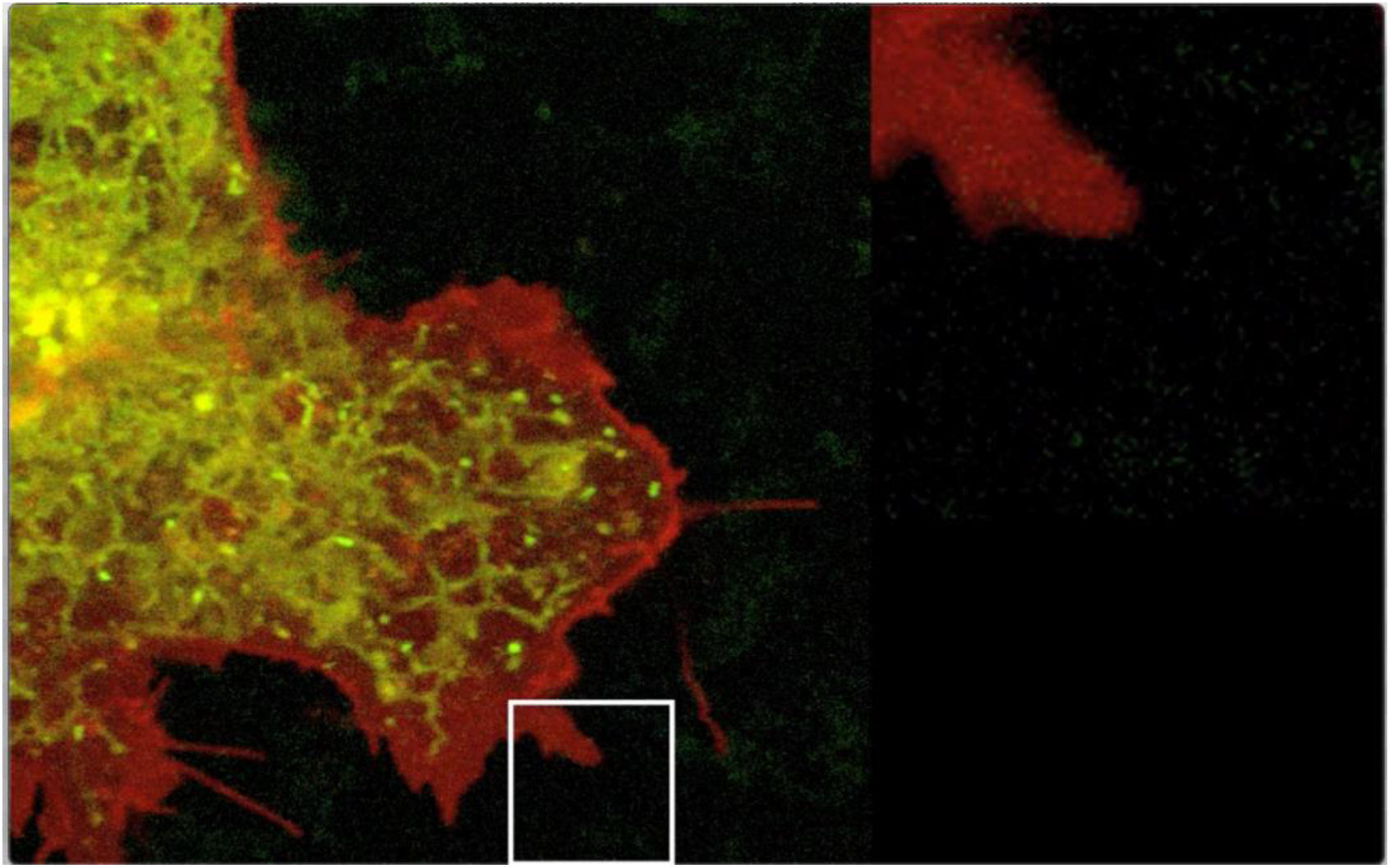
CPE-mCherry secretion in a living cell. Movie showing a living cell co-expressing CPE-GFP (green) with the PM marker lipid anchored mCherry (red)

**Movie S2.**
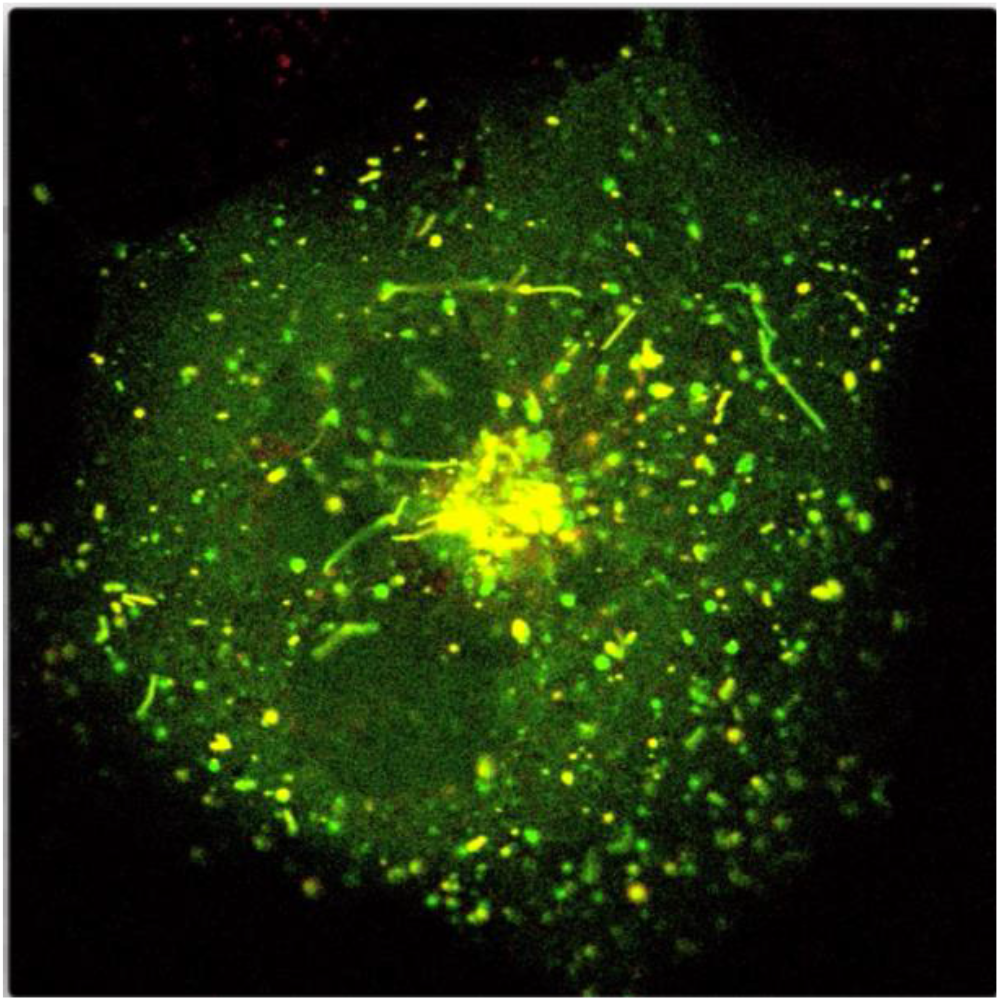
CPE-mCherry localization in lysosomes in living cells. Movie showing a living cell co-expressing CPE-mCherry (red) with the lysosomal marker LGP120-GFP (green)

## REFERENCES

Andrei C, Dazzi C, Lotti L, Torrisi MR, Chimini G, Rubartelli A (1999) The secretory route of the leaderless protein interleukin 1beta involves exocytosis of endolysosome-related vesicles. Mol Biol Cell 10: 1463–1475

Arnaoutova I, Jackson CL, Al-Awar OS, Donaldson JG, Loh YP (2003) Recycling of Raft-associated prohormone sorting receptor carboxypeptidase E requires interaction with ARF6. Mol Biol Cell 14: 4448–4457

Ashkenazy H, Abadi S, Martz E, Chay O, Mayrose I, Pupko T, Ben-Tal N (2016) ConSurf 2016: an improved methodology to estimate and visualize evolutionary conservation in macromolecules. Nucleic Acids Res 44: W344–350

Bennett JP, Cockcroft S, Gomperts BD (1979) Ionomycin stimulates mast cell histamine secretion by forming a lipid-soluble calcium complex. Nature 282: 851–853

Brakch N, Rist B, Beck-Sickinger AG, Goenaga J, Wittek R, Burger E, Brunner HR, Grouzmann E (1997) Role of prohormone convertases in pro-neuropeptide Y processing: coexpression and in vitro kinetic investigations. Biochemistry 36: 16309–16320

Bright NA, Reaves BJ, Mullock BM, Luzio JP (1997) Dense core lysosomes can fuse with late endosomes and are re-formed from the resultant hybrid organelles. J Cell Sci 110 (Pt 17): 2027–2040

Carvalho MH, Prat JC, Garcia AG, Kirpekar SM (1982) Ionomycin stimulates secretion of catecholamines from cat adrenal gland and spleen. Am J Physiol 242: E137–145

Cheng Y, Cawley NX, Loh YP (2013) Carboxypeptidase E/NFalpha1: a new neurotrophic factor against oxidative stress-induced apoptotic cell death mediated by ERK and PI3-K/AKT pathways. PLoS One 8: e71578

Corrigall VM, Bodman-Smith MD, Brunst M, Cornell H, Panayi GS (2004) Inhibition of antigen-presenting cell function and stimulation of human peripheral blood mononuclear cells to express an antiinflammatory cytokine profile by the stress protein BiP: relevance to the treatment of inflammatory arthritis. Arthritis Rheum 50: 1164–1171

Cranfill PJ, Sell BR, Baird MA, Allen JR, Lavagnino Z, de Gruiter HM, Kremers GJ, Davidson MW, Ustione A, Piston DW (2016) Quantitative assessment of fluorescent proteins. Nat Methods 13: 557–562

Das B, Sabban EL, Kilbourne EJ, Fricker LD (1992) Regulation of carboxypeptidase E by membrane depolarization in PC12 pheochromocytoma cells: comparison with mRNAs encoding other peptide- and catecholamine-biosynthetic enzymes. J Neurochem 59: 2263–2270

Dayel MJ, Hom EF, Verkman AS (1999) Diffusion of green fluorescent protein in the aqueous-phase lumen of endoplasmic reticulum. Biophys J 76: 2843–2851

Delpino A, Castelli M (2002) The 78 kDa glucose-regulated protein (GRP78/BIP) is expressed on the cell membrane, is released into cell culture medium and is also present in human peripheral circulation. Biosci Rep 22: 407–420

Dhanvantari S, Loh YP (2000) Lipid raft association of carboxypeptidase E is necessary for its function as a regulated secretory pathway sorting receptor. J Biol Chem 275: 29887–29893

Dikeakos JD, Reudelhuber TL (2007) Sending proteins to dense core secretory granules: still a lot to sort out. The Journal of cell biology 177: 191–196

Fedele AO, Proud CG (2020) Chloroquine and bafilomycin A mimic lysosomal storage disorders and impair mTORC1 signalling. Biosci Rep 40

Fricker LD (1988) Carboxypeptidase E. Annu Rev Physiol 50: 309–321

Fricker LD, Snyder SH (1983) Purification and characterization of enkephalin convertase, an enkephalin-synthesizing carboxypeptidase. J Biol Chem 258: 10950–10955

Fukuda M (1991) Lysosomal membrane glycoproteins. Structure, biosynthesis, and intracellular trafficking. J Biol Chem 266: 21327–21330

Fukuda M (2008) Regulation of secretory vesicle traffic by Rab small GTPases. Cell Mol Life Sci 65: 2801–2813

Ghosh S, Dellibovi-Ragheb TA, Kerviel A, Pak E, Qiu Q, Fisher M, Takvorian PM, Bleck C, Hsu VW, Fehr AR et al (2020) beta-Coronaviruses Use Lysosomes for Egress Instead of the Biosynthetic Secretory Pathway. Cell 183: 1520–1535 e1514

Hamano T, Gendron TF, Causevic E, Yen SH, Lin WL, Isidoro C, Deture M, Ko LW (2008) Autophagic-lysosomal perturbation enhances tau aggregation in transfectants with induced wild-type tau expression. Eur J Neurosci 27: 1119–1130

Hurtley SM, Bole DG, Hoover-Litty H, Helenius A, Copeland CS (1989) Interactions of misfolded influenza virus hemagglutinin with binding protein (BiP). J Cell Biol 108: 2117–2126

Ji L, Wu HT, Qin XY, Lan R (2017) Dissecting carboxypeptidase E: properties, functions and pathophysiological roles in disease. Endocr Connect 6: R18–R38

Kern J, Untergasser G, Zenzmaier C, Sarg B, Gastl G, Gunsilius E, Steurer M (2009) GRP-78 secreted by tumor cells blocks the antiangiogenic activity of bortezomib. Blood 114: 3960–3967

Kim T, Gondre-Lewis MC, Arnaoutova I, Loh YP (2006) Dense-core secretory granule biogenesis. Physiology (Bethesda) 21: 124–133

Kneen M, Farinas J, Li Y, Verkman AS (1998) Green fluorescent protein as a noninvasive intracellular pH indicator. Biophys J 74: 1591–1599

Laulagnier K, Schieber NL, Maritzen T, Haucke V, Parton RG, Gruenberg J (2011) Role of AP1 and Gadkin in the traffic of secretory endo-lysosomes. Mol Biol Cell 22: 2068–2082

Li Z, Zhuang M, Zhang L, Zheng X, Yang P, Li Z (2016) Acetylation modification regulates GRP78 secretion in colon cancer cells. Sci Rep 6: 30406

Ling H, Ardjomand P, Samvakas S, Simm A, Busch GL, Lang F, Sebekova K, Heidland A (1998) Mesangial cell hypertrophy induced by NH4Cl: role of depressed activities of cathepsins due to elevated lysosomal pH. Kidney Int 53: 1706–1712

Luo DX, Huang MC, Ma J, Gao Z, Liao DF, Cao D (2011) Aldo-keto reductase family 1, member B10 is secreted through a lysosome-mediated non-classical pathway. Biochem J 438: 71–80

Manser E, Fernandez D, Lim L (1991) Processing and secretion of human carboxypeptidase E by C6 glioma cells. Biochem J 280 (Pt 3): 695–701

Marin-Briggiler CI, Gonzalez-Echeverria MF, Munuce MJ, Ghersevich S, Caille AM, Hellman U, Corrigall VM, Vazquez-Levin MH (2010) Glucose-regulated protein 78 (Grp78/BiP) is secreted by human oviduct epithelial cells and the recombinant protein modulates sperm-zona pellucida binding. Fertil Steril 93: 1574–1584

Mauvezin C, Nagy P, Juhasz G, Neufeld TP (2015) Autophagosome-lysosome fusion is independent of V-ATPase-mediated acidification. Nat Commun 6: 7007

McGirr R, Guizzetti L, Dhanvantari S (2013) The sorting of proglucagon to secretory granules is mediated by carboxypeptidase E and intrinsic sorting signals. J Endocrinol 217: 229–240

Niu TK, Pfeifer AC, Lippincott-Schwartz J, Jackson CL (2005) Dynamics of GBF1, a Brefeldin A-sensitive Arf1 exchange factor at the Golgi. Mol Biol Cell 16: 1213–1222

Patterson GH, Lippincott-Schwartz J (2002) A photoactivatable GFP for selective photolabeling of proteins and cells. Science 297: 1873–1877

Pols MS, Klumperman J (2009) Trafficking and function of the tetraspanin CD63. Exp Cell Res 315: 1584–1592

Ramamoorthy P, Whim MD (2008) Trafficking and fusion of neuropeptide Y-containing dense-core granules in astrocytes. J Neurosci 28: 13815–13827

Saito N, Takeuchi T, Kawano A, Hosaka M, Hou N, Torii S (2011) Luminal interaction of phogrin with carboxypeptidase E for effective targeting to secretory granules. Traffic 12: 499–506

Shinoda H, Shannon M, Nagai T (2018) Fluorescent Proteins for Investigating Biological Events in Acidic Environments. Int J Mol Sci 19

Skalka N, Caspi M, Lahav-Ariel L, Loh YP, Hirschberg K, Rosin-Arbesfeld R (2016) Carboxypeptidase E (CPE) inhibits the secretion and activity of Wnt3a. Oncogene 35: 6416–6428

Tang SS, Zhang JH, Liu HX, Li HZ (2009) PC2/CPE-mediated pro-protein processing in tumor cells and its differentiated cells or tissues. Mol Cell Endocrinol 303: 43–49

Tapper H, Sundler R (1990) Role of lysosomal and cytosolic pH in the regulation of macrophage lysosomal enzyme secretion. Biochem J 272: 407–414

Topalidou I, Cattin-Ortola J, Hummer B, Asensio CS, Ailion M (2020) EIPR1 controls dense-core vesicle cargo retention and EARP complex localization in insulin-secreting cells. Mol Biol Cell 31: 59–79

Varlamov O, Fricker LD (1996) The C-terminal region of carboxypeptidase E involved in membrane binding is distinct from the region involved with intracellular routing. J Biol Chem 271: 6077–6083

Villeneuve J, Bassaganyas L, Lepreux S, Chiritoiu M, Costet P, Ripoche J, Malhotra V, Schekman R (2018) Unconventional secretion of FABP4 by endosomes and secretory lysosomes. J Cell Biol 217: 649–665

Yaffe Y, Hugger I, Yassaf IN, Shepshelovitch J, Sklan EH, Elkabetz Y, Yeheskel A, Pasmanik-Chor M, Benzing C, Macmillan A et al (2015) The myelin proteolipid plasmolipin forms oligomers and induces liquid-ordered membranes in the Golgi complex. J Cell Sci 128: 2293–2302

